# Sex tweaks eye color

**DOI:** 10.1101/2024.07.26.605233

**Authors:** Paola Bressan

## Abstract

Here I challenge the conventional view of eye-color inheritance by showing that eye color depends (doubly) on sex: the sex of the parent and the sex of the child. I rely on the eye color records of over 30,000 Italians—a single-ethnicity population that features the largest genetic and eye-color diversity in Europe. First, more men than women express eye colors at the two extremes of the melanin range (blue and dark-brown) and more women than men express colors in the middle (green, hazel, brown, and mixed). Second, dark-brown-eyed mothers produce more *sons* with dark-brown eyes and fewer with very-light-brown ones than do dark-brown-eyed fathers. Symmetrically, dark-brown-eyed fathers produce more *daughters* with dark-brown eyes and fewer with very-light-brown ones than do dark-brown-eyed mothers. All of this suggests that human eye pigmentation is controlled by a gene sitting on chromosome X—a chromosome which fathers pass on only to daughters, and sons inherit only from mothers. Such knowledge can improve forensic tools’ ability to predict eye color from DNA found at crime scenes and help in the identification of a known individual’s unknown birth parents. On a grander scale, these findings cast a new light on the origin of other, more fateful sex differences.

---

> “Even the iris of the eye is sometimes more brightly-coloured in the male than in the female.”
>
> — Charles Darwin, *The Descent of Man and Selection in Relation to Sex, Chapter XIII*

Blue eyes arise from a small genetic fluke, a mutation which expresses itself only when we inherit it from both parents. Or so the story used to go. As it now turns out, blue eyes can show up on people who do *not* carry this mutation (Pośpiech et al., 2016; Martinez-Cadenas et al., 2013); for mysterious reasons, all of them appear to be males (see Figure 3 in Pośpiech et al., 2016). Every bit as oddly, women tend to have darker eyes than men bearing the same genotype. They are less often blue-eyed, and more often green- or brown-eyed, than are men; although whether any such differences are statistically significant, or show up at all, varies a great deal across samples (Martinez-Cadenas et al., 2014; Sulem et al., 2007; Pietroni et al., 2014; Liu et al., 2010; Liu et al., 2014; Lock-Andersen et al., 1998; Grant & Lauderdale, 2002).

The mutation that produces blue eyes is a one-letter alteration of a short sequence on the gene *HERC2* (Sturm et al., 2008). This sequence, rs12913832, is highly conserved across species and regulates the ability of the nearby gene *OCA2* to produce melanin, the pigment that darkens eyes, hair, and skin. The ancestral allele at rs12913832 stimulates melanin production, leading to dark-brown eyes. The mutated allele reduces it fivefold (Duffy, 2015), allowing the eyes to look blue—or, presumably depending on lesser-known genes and modifiers, grey, green, amber, hazel, and all manner of mixtures.

From a DNA sample, indeed, forensic eye-color prediction systems such as IrisPlex can estimate whether someone’s eyes are likely to be blue, brown or intermediate (Walsh et al., 2011; Walsh et al., 2012). And yet these models’ accuracy, while exceeding 90% for blue and brown (almost entirely thanks to the *HERC2* blue/brown allele Pietroni et al., 2014), drops precipitously for intermediate colors. Incorporating sex into IrisPlex has been attempted more than once, but with inexplicably inconsistent results (Martinez-Cadenas et al., 2014; Pietroni et al., 2014; Liu et al., 2010; Liu et al., 2014).

Here I put forth a simple solution to all these mysteries and test some surprising, counterintuitive predictions of my proposal on a very large new dataset.

## The hypothesis: Eye color is linked to the X chromosome

The *HERC2* blue/brown switch that regulates the *OCA2* gene—whose malfunctioning produces the form of albinism in which eyes, hair and skin carry very little pigment, or none—resides on our chromosome 15. Sex does not come into the picture here, for chromosome 15 is inherited identically by sons and daughters—as are all other chromosomes that host known markers of iris color. And yet, an even more common type of albinism proves that eye melanin is heavily influenced by one gene sitting on chromosome X, which is of course strongly related to sex. This gene, *OA1*, is implicated in the growth of melanosomes, the cellular structures that manufacture and store melanin. Most of this gene’s mutations cause ocular albinism type 1, which affects only the eyes (hair and skin colors remain normal) and is inherited in the same manner as are daltonism, hemophilia, and other X-linked recessive diseases: mothers pass it on and sons express it.

I suggest that a mutation in a regulatory element of this (or another such X-linked) gene reduces iris pigmentation relative to the original allele, which increases it. As I will show, the presence of both the mutated and ancestral alleles in a population would engender precisely the pattern of sex differences in eye color (along with prediction errors and apparent contradictions) one observes. And if you bear with me till the end, you may concur that this is far from a small matter of lighter or darker shades. Indeed, the part played in the evolution of human eye color by this unrecognized X-linked mutation must have been a portentous one.

### Testing the hypothesis on 30,000 fresh pairs of eyes

I now lay out the hypothesis’ main implications and test them on a new dataset of over 30,000 individuals (10,344 participants, their parents, and current partners). These data, which we have collected from 2015 to 2022, reap the unique benefit of having been gathered in Italy: that is, on a single-ethnicity population that features the largest genetic diversity in Europe (Raveane et al., 2019), and in which the entire range of eye colors is far better represented than further north or further south (Salvoro et al., 2019).

The dataset collates the data of all eye-color studies we have done to date. Participants were recruited via online social media. In all studies, people were asked to report their own eye color, the eye color of their parents, and the eye color of their partner if they had one. Options were: “dark brown”, “brown”, “hazelnut (very light brown)”, “green”, “grey”, “blue”, and “other” (with the invitation to specify the exact color). The data of participants who reported being adopted, not being Italian, having at least one non-Italian parent, or having previously participated in a similar survey on eye color were excluded.

The analyses presented here include all individuals whose eye color fell into the following predefined categories: blue, green, hazelnut (very light brown), brown, dark brown. For nonparametric correlations, these were coded on a 5-point ordinal scale from light to dark (as in Bressan, 2020), reflecting the continuum in the amount of melanin in the iris. Incidentally, the results did not change (see supplementary material) using a looser, 4-point scale which included grey and “other” eye colors, as follows: Light (blue, grey, blue/grey combinations), Intermediate (green, all manner of mixtures containing green, very light brown), Dark (brown), Very Dark (dark brown, black). The final sample for the analyses reported below comprises 32,776 people (a younger cohort of 14,436 participants and partners plus an older cohort of 18,330 mothers and fathers). Note that here the eye-color records of parents and children, who are not genetically independent, are never collapsed together; eye-color frequency distributions always portray unrelated individuals.

### Eye color depends on sex

Men, who are stuck with a single X chromosome, are bound to express the single allele this bears: either the ancestral one, which codes for extra melanin, or the mutated, faulty variant, which does not. The mutated allele allows the iris to look blue if this is the color dictated by autosomal genes (i.e., those not on sex chromosomes). The ancestral allele, by increasing pigmentation, raises the odds of dark-brown eyes in those carriers of one *HERC2* “brown” allele who would otherwise showcase a lighter shade of brown. Women, on the other hand, carry *two* X chromosomes and randomly inactivate either one in different cells. Whenever one of the two X-linked alleles is mutated (in technical jargon, “derived”) and the other ancestral, then, female irides will express a mosaic of both and so display intermediate amounts of melanin. Hence, women will feature a lower frequency of eye colors at the extremes of the melanin range, such as blue and dark brown, than do men; and a higher frequency of intermediate and mixed eye colors, such as green, lighter shades of brown, blue-green or green-brown (Figure 1).

**Figure 1.**
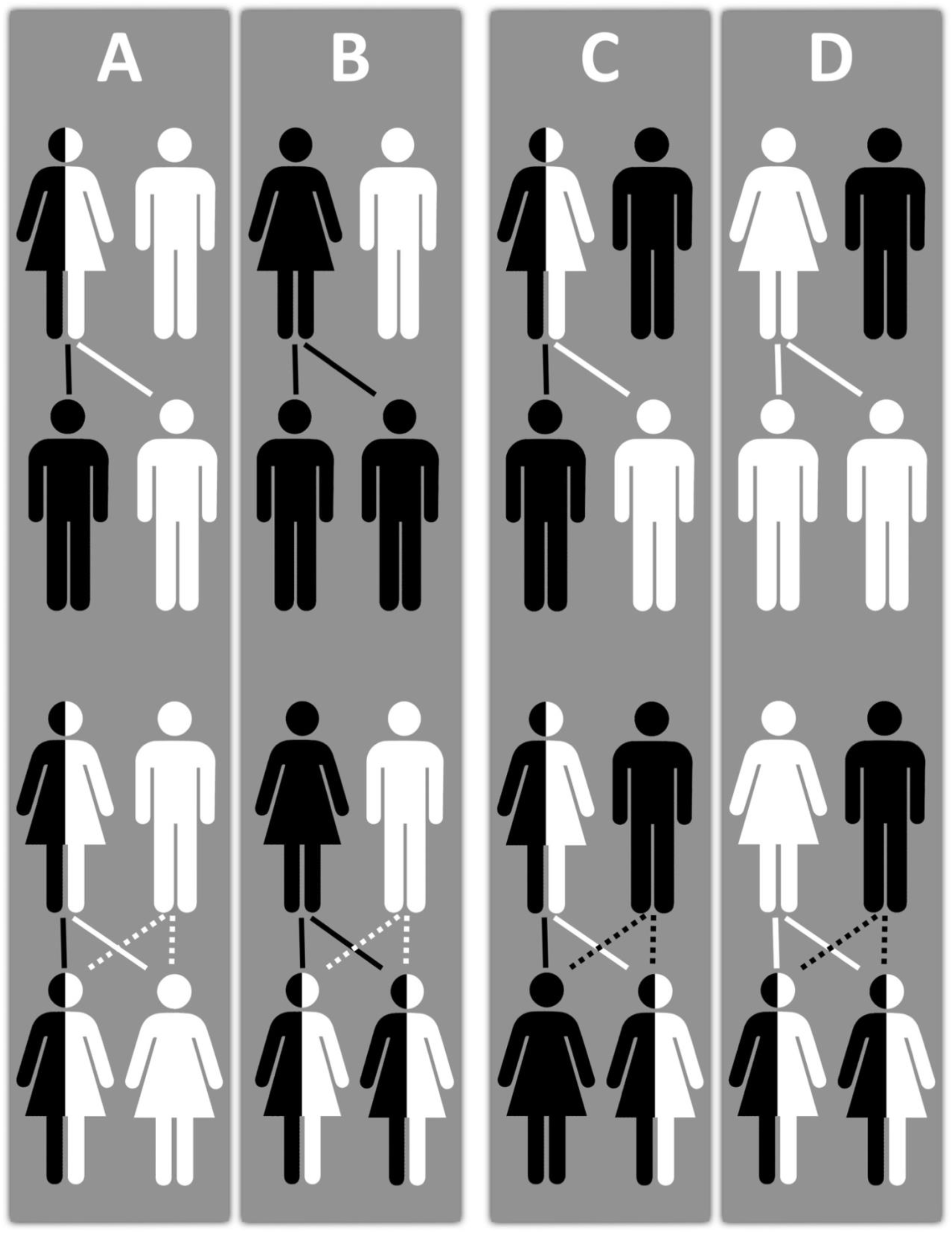
Offspring’s probability of inheriting the X-linked ancestral, iris-darkening (in black) and the derived, iris-lightening (in white) alleles, when the two parents bear different X-linked alleles. Sons **(top)** carry a single X, inherited from their mother, and thus fully express either the ancestral or the derived allele. Daughters **(bottom)** carry two Xs, and normally express both alleles. Therefore, whichever the eye color dictated by autosomal genes, more men than women will feature very light or very dark irides (**top:** all sons are either in black or in white), and more women than men will feature irides of intermediate lightness (**bottom:** most daughters are half in black and half in white). **Panels B, C, D:** Offspring’s probability of carrying either X-linked allele when one parent has dark-brown eyes, hence bears only the ancestral X-linked allele(s), and the other has lighter eyes (blue, green, very light brown, or brown), hence bears at least one copy of the derived X-linked allele. **Top:** sons of dark-brown-eyed mothers are far *more* likely to be dark-eyed (**B:** 100% of sons are in black) than sons of dark-brown-eyed fathers (**C-D:** 25% of sons are in black). **Bottom:** daughters of dark-brown-eyed mothers are slightly *less* likely to be dark-eyed (**B:** 0% of daughters are in black) than daughters of dark-brown-eyed fathers (**C-D:** 25% of daughters are in black).

All these sex differences must show up especially in carriers of at least one *HERC2* “blue” allele, for whom any extra melanin will produce noticeable iris darkening. If they are female, *HERC2* blue/blue homozygotes will be less likely to be blue-eyed, and *HERC2* blue/brown heterozygotes will be more likely to be brown-eyed. Incidentally this also implies that, relative to men with the same genotype, female blue/blue homozygotes (who, owing to the extra melanin, are pulled away from blue, towards the full gamut of colors) may show a larger *variability* in eye color; and female blue/brown heterozygotes (who, owing to the extra melanin, are pushed towards brown, away from the full gamut of colors) a smaller one. This appears to be the case: see Figure 1 in Pietroni et al., 2014, and Figure 3 in Martinez-Cadenas et al., 2013.

The sex biases in eye-color frequency just described are very clear in my data (Figure 2) and perfectly match the IrisPlex prediction errors, whether the latter are recognized as due to some unexplained sex difference (Martinez-Cadenas et al., 2013; Pietroni et al., 2014), or not (Andersen et al., 2013).

**Figure 2.**
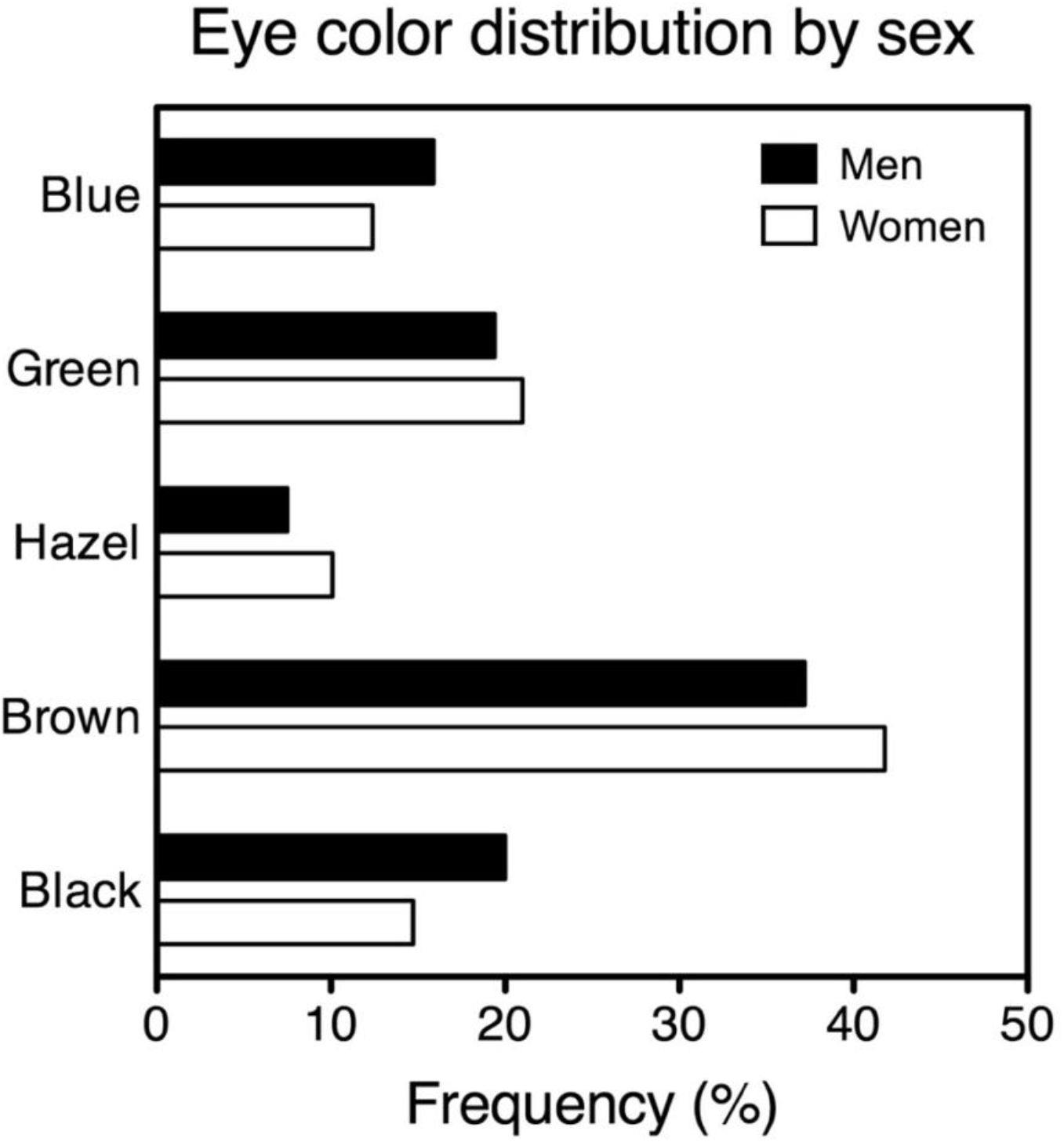
Eye color distribution in a sample of 18,330 Italians (9,026 men, 9,304 women). More men than women have blue or dark-brown (“black”) eyes; more women than men have green, very-light-brown (“hazel”), or brown eyes, *χ*^*2*^ (4)=180.1, p<.0001. All sex differences are separately significant (see supplementary material for contingencies tables and statistics). Only the participants’ parents were included in this sample, reducing noise in two ways. First, the numbers of women (mothers) and men (fathers) that are compared are balanced, even though we had many more female than male participants. Second, the female and male eye colors that are compared were assessed by the same person (the child), which avoids any confounds due to one sex being potentially more likely than the other to give detailed and nuanced descriptions of their own eye color.

Note that sex differences are driven entirely by women who possess *both* an ancestral and a derived X-linked allele. Women bearing two identical alleles on their two Xs (one of which is switched off) are just like men bearing that allele’s single copy on their single X. The more evenly distributed the two alleles are in a population, the higher the likelihood that any woman is carrying both of them, and the larger the sex effect. At the same time, as pointed out above, the effect also increases with the relative number of *HERC2* blue-allele carriers. The interplay between these two ratios (*HERC2* derived vs ancestral, X-linked derived vs ancestral) introduces all manner of variability. For example, if the X-linked ancestral allele is fixed, *HERC2* blue and brown alleles might primarily modulate variations in brown iris color (with two blue alleles making brown eyes lighter and greener, but not blue)—as indeed they do in southern Asians (Edwards et al., 2016). If the two X-linked alleles are optimally balanced, instead, one ought to expect an assortment of light and dark blues and greys and greens wherever *HERC2* blue alleles abound, of brown shades wherever they are in short supply. Yet if all brands of brown are fitted into the same “brown” slot, the X contribution will remain largely invisible. Alas, very few studies (with the notable exception of Edwards et al., 2016) have paid any attention to the multiplicity of browns, in or out of Europe. All of the above implies that the sex effect will not only differ in nature and size between populations, but also be easier or harder to find depending on which eye color categories one is using.

Thus, it is only to be expected that the extent of male-female differences will vary even vastly and in a seemingly haphazard manner, especially when one compares samples that are relatively small and/or come from distinct populations. As a case in point, the male advantage found so far for blue eyes across seven independent samples ranges from 18% to a nonsignificant -1% (Martinez-Cadenas et al., 2014). In my two Italian samples it is 3% (younger cohort: male and female participants vs their current opposite-sex partners) and 4% (older cohort: participants’ fathers vs mothers). Please note that any assortative mating for eye color potentially occurring in these pairs (i.e., a tendency to choose partners similar to oneself) would if anything reduce, never inflate, sex differences.

### Sons take after mothers, and daughters after fathers

A compelling prediction of the idea that eye color is X-linked can be tested by looking at what happens when one parent’s eyes are dark brown and the other’s are not (Figure 1, panels B, C, D). The dark-brown-eyed parent is expected to bear one (if he is a father) or two (if she is a mother) copies of the *ancestral*, melanin-boosting X-linked allele, because his or her irides would be a lighter shade of brown or some other color otherwise. Yet the other parent, being lighter-eyed, must carry exactly one (if he is a father) or at least one (if she is a mother) copy of the *derived* allele.

Now, men invariably inherit their sole X chromosome from their mother, as their father provides the Y. When it comes to X-linked genes, then, sons are entirely in thrall to their mothers whilst fathers count for nothing. This implies that sons of dark-brown-eyed mothers should be more likely to have dark-brown eyes, and less likely to have very-light-brown eyes, than sons of dark-brown-eyed fathers (Figure 1, top: compare panel B with panels C-D). This turns out to be true (Figure 3, left panel).

**Figure 3.**
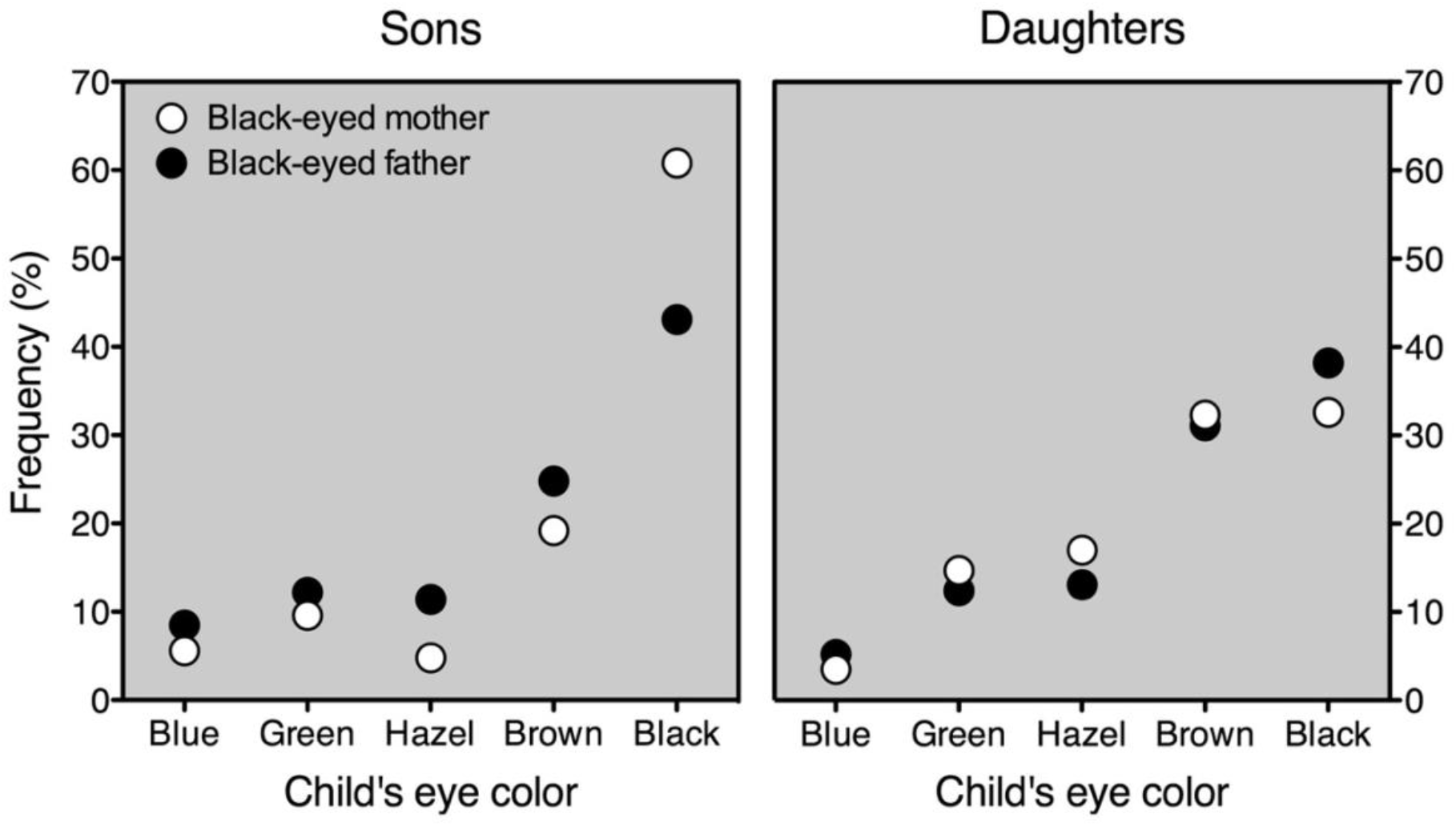
Offspring eye color frequencies when only the mother (open symbols) or only the father (closed symbols) has dark-brown eyes (here labelled as “black”). Left panel: sons (*N*=371). Sons of dark-brown-eyed mothers are significantly more likely to have dark-brown eyes, and less likely to have very-light-brown ones (here labelled as “hazel”), than sons of dark-brown-eyed fathers: open symbols are higher than closed ones for “black”, lower for “hazel”. Right panel: daughters (*N*=1634). Daughters of dark-brown-eyed fathers are slightly but significantly more likely to have dark-brown eyes, and less likely to have very-light-brown ones, than daughters of dark-brown-eyed mothers: closed symbols are higher than open ones for “black”, lower for “hazel”. See supplementary material for contingencies tables and statistics.

Things go elsewise for daughters, who acquire one X from their mother and the other from their father. The eye-color data mirror this overall balance in parental contribution (Figure 3, right panel). Nonetheless, the asymmetrical nature of X chromosome expression and inheritance ought to create a small paternal bias. All daughters of a dark-brown-eyed father carry the ancestral allele expressed by him (Figure 1C-D, bottom), but not all possess the derived allele that is partly or fully expressed by their lighter-eyed mother. Alongside the derived allele, indeed, their mother could be bearing a copy of the ancestral one—which she will proceed to hand down to half her daughters (Figure 1C, bottom). On the other hand, all daughters of a dark-brown-eyed mother carry exactly one copy of the two ancestral alleles expressed by their mother *and* one copy of the derived allele expressed by their lighter-eyed father (Figure 1B, bottom). And in fact, daughters of dark-brown-eyed fathers have slightly but significantly higher odds of dark-brown eyes, and slightly but significantly lower odds of very-light-brown eyes, than daughters of dark-brown-eyed mothers (Figure 3, right panel).

These findings reflect a more general, rather striking sex parallelism. In eye color, sons resemble their mothers more than their fathers, *rs*(1595)=.40 vs *rs*(1562)=.35, *z*=1.7, p=.048, one-tailed, and daughters their fathers more than their mothers, *rs*(6827)=.40 vs *rs*(7036)=.35, *z*=3.3, p=.0004, one-tailed. This double-sided parent-of-origin effect, which is extraordinarily hard to explain any other way, supports quite forcefully the idea that human eye color is X-linked.

### Sex and the IrisPlex

The data presented here suggest that the disappointing effects of sex as a predictor in IrisPlex arise from “femaleness” being treated as a risk factor for darker eyes. True, this rule is bound to sharpen the model’s performance in all cases where being female does darken the eyes. These are the genotypes that produce nonbrown-eyed males—that is, *HERC2* “blue/blue” homozygotes and a portion of “blue/brown” heterozygotes. But any such refinements in predictive power will be partly or fully offset, depending on the sample, by all cases where being female *lightens* the eyes. These are the genotypes that produce the brown range of irides—*HERC2* “brown/brown” homozygotes and a sizeable portion of “blue/brown” heterozygotes. For example, female heterozygotes carrying an X-linked derived allele will have their autosomally brown eyes (“brown”) lighten into very light brown or hazel (“intermediate”), and yet the sex-savvy model will erroneously predict the opposite change, from “intermediate” to “brown”.

In general, it is easy to see why most of the IrisPlex prediction failures occur for intermediate eye colors. Males will be pulled away from the “intermediate” category in either direction (green irides lightening into blue, hazel irides darkening into dark-brown) depending on whether their single X-linked allele is, respectively, derived or ancestral. Females will be pulled *away* from “intermediate”, just like males, if they carry two identical copies of the X-linked gene; but pushed *towards* “intermediate” (blue irides darkening into green, dark-brown irides lightening into hazel) if they carry two different copies of it.

### The many shades of Brown

Sex differences in eye color, then, can be attributed to the averaging effect of X-inactivation in females. Crucial to figuring this out was the present discovery that more men than women have dark-brown eyes. This piece of knowledge would have been harder to come across before, simply because in all previous studies brown and dark brown were lumped together into just one “brown” category. Such a classification practice is actually exacerbated by recent, more sophisticated measures of eye color, such as the PIE-score. This expresses the ratio between blue and brown pixels in a digital image of the iris, and hence makes eye color “objective” and “quantitative”. Still, it can currently differentiate neither between light and dark blue nor, more momentously, between light and dark brown (Andersen et al., 2013). And yet, only this distinction does permit one to properly use sex as a predictor.

It will not have escaped the reader, by the way, that the work presented here could help models such as IrisPlex identify the unknown parents of a known individual by assessing the probability that they have a given eye color. For example, a dark-brown-eyed male will be disproportionately likely, and a very-light-brown-eyed male unlikely, to have been born from a dark-brown-eyed mother.

### Males in the tails

The higher proportion of males at the low and high ends of the iris melanin distribution inevitably reminds one of many other polygenic traits with more men in the tails and more women in the middle. These include birth weight, adult height and weight, running speed, and university grades (Lehre et al., 2009). In some such characteristics, as male height, the X-chromosome is demonstrably on board (Pan et al., 2007; Reinhold & Engqvist, 2013). In other traits, particularly those concerning the mind, the involvement of X-linked genes has been advocated (e.g., Johnson et al., 2009; Skuse, 2006) and disputed (e.g., Turkheimer & Halpern, 2009) just as passionately. All manner of hormonal or evolutionary or sociocultural explanations have been proposed instead (see Lehre et al., 2009).

The parent-of-origin findings presented here do not admit any such alternative explanations. Indeed, that this sort of results is scarcely open to the usual multitude of nongenetic interpretations encourages one to test whether sons may resemble mothers more than fathers, and daughters resemble fathers more than mothers, in other heritable traits whose distribution depends on sex. That children may take after their opposite-sex parent in some cognitive abilities (see Giummo & Johnson, 2012), for example, is a claim that psychological or physiological theories of any description would regard as outlandish. Thus, this sort of finding—that is bound to support X chromosome’s involvement whichever way one slices it—would greatly upset such schools of thought, or at least any pretence that their approaches could lead to complete explanations of behaviour. As we shall now see, the most promising traits one may wish to put to the test are those that are likely to have been shaped, or at least fine-tuned, by sexual selection: that is, by mate choice.

### The far view: Sex and the X

That eye color is partly controlled by the X chromosome makes a great deal of sense in the light of evolution. The X harbours an inordinate proportion of genes that control sexually selected traits (Reinhold, 1998). Blue eyes are such a trait: in a prospective partner, people of European ancestry prefer light, colorful (above all, blue) eyes over dark-brown ones. This is suggested by preference data collected in Italy (Bressan & Damian, 2018; Bressan, 2020; Bressan, 2024) but most compellingly by the swift, massive diffusion across Europe—well before skin lightening—of the *HERC2* mutation that made light eyes possible (Bressan, 2024). It looks like this variant, starting from a sole copy (Eiberg et al., 2008), had already swept to fixation in hunter-gatherers inhabiting northern Europe 8,000 years ago: every single specimen uncovered so far has turned out to have been blue-eyed (Mathieson et al., 2015). Today, in times of heavier gene flow between populations, the mutation is present in an impressive 70% to 95% of northern Europeans and in substantial proportions of southern Europeans, southwest Asians, and northern Africans; up to 20% of central and southern Asians carry it as well (Kidd et al., 2020).

I have made the case that the steep ascent of human blue eyes came about via runaway evolution, the self-reinforcing coevolution of a trait and the preference for it (Bressan, 2024). In the same spirit, any mutation (on whichever chromosome) that happens to reduce melanin in the iris would spread fast, insofar as it produces brighter, more colorful, sexier eyes in its carriers. In a sunlit world, however, poorly pigmented irides are a handicap (Bressan, 2024), and a sexually selected trait must reap a reproductive benefit larger than its cost to make it to the next generation. The cost of showcasing blue eyes is the same to males and females but the benefit looms larger for males, because, in humans (Todd et al., 2007) as in most every animal species, it is mainly the females who choose males and not the other way around. Therefore, males are under heavier pressure to express their recessive, sexually attractive, costly traits than are females—and recessive alleles residing on the X chromosome are always expressed in male carriers. Indeed, an X-linked recessive gene that favours males will spread even if the cost to females exceeds the gain to males (Rice, 1984). In broader terms, one expects a trait to become linked to the X whenever it is expressed in both sexes but its optimal size differ between them (Reinhold, 1998). This is precisely the case with blue eyes, whose smaller benefit/cost ratio for females than for males brings about a mild form of such so-called sexually antagonistic selection. X linkage of an eye-lightening allele provides a splendidly economical solution by ensuring, at the same time, bluer, sexier irides in male carriers and browner, safer irides in female ones.

Here comes a final twist. Because by boosting melanin it darkens the iris, the X-linked ancestral allele is incompatible with blue eyes. As long as such a gene remains in its original form, a *HERC2* “blue” mutant would feature brown irides at best—not yet the stuff of seduction, hardly a spur for any spectacular brand of runaway sexual selection. Thus the X-linked derived allele, whether or not it appeared on the scene first, must have acted as a *prerequisite* for the *HERC2* variant to manifest itself in flying colors. One gets the idea indeed that the much acclaimed blue-eye mutation owes its triumph to this obscure, unnamed, unsung change on our X chromosome.

### Implications and applications

Drawing on a database of over 30,000 Italians—a population that features the largest genetic and eye-color diversity in Europe—I have presented empirical and logical evidence that human eye pigmentation depends on sex in ways that imply the contribution of an X-linked gene. Such a conclusion has implications for people from various walks of science and life.

1. Genetics: these findings encourage researchers to look at the X chromosome for an eye-color gene that virtually all dark-brown-eyed males carry in its ancestral form and virtually all blue-eyed males in its mutated, loss-of-function form.
2. Evolutionary biology: these findings (a) suggest that the X-linked eye-lightening mutation preceded the full expression of the *HERC2* one, and (b) lend support to the theory (Bressan, 2024) that human blue eyes evolved as costly sexual ornaments whose net advantage was, as is typical for such traits, larger in the more competitive sex (males) than in the choosier one (females).
3. Ecology and epidemiology: these findings (a) clarify why sex differences in eye color, depending as they do on the frequency of both the X-linked and *HERC2* mutations in the population, would not be expected to turn up in every European sample, and hence (b) make plain that any past failures to replicate the effect by no means signify this is spurious or unreliable.
4. Forensic methods: these findings (a) prove the importance of developing iris color measures that distinguish between shades of brown, and (b) entail that sex has fared poorly as an eye-color predictor in IrisPlex owing not to true lack of effects but to mistaken assumptions about what these should be.
5. Forensic applications: these findings (a) specify how sex sways eye color between and within the categories of blue, green, and brown, and hence (b) can improve the accuracy of current and future models in predicting eye color from DNA collected at crime scenes.
6. Genealogy: these findings (a) define how the sex of the parent and the sex of the child alter the probability that they feature certain eye color combinations, and hence (b) can help in the identification of a known individual’s unknown birth parents.
7. The public at large: these findings (a) spell out a mechanism that allows for blue eyes in people who do not carry two copies of the recessive *HERC2* blue-eye allele, and hence (b) show that blue eyes in the offspring of a brown-eyed father need not arouse doubts about paternity.

## Data accessibility

All data, analysis scripts, and annotated outputs are publicly available as supplementary material via the Open Science Framework and can be accessed at https://osf.io/eaknc [https://bit.ly/3Sq3Kf9]

## Acknowledgements

I am, as usual, much indebted to Peter Kramer for his encouragement and invaluable comments. I am grateful to all the students who collected data over the years: Valeria Damian (2015-16), Isabella Ragona (2016-17), Valentina Mini (2017-18), Paola Monegatti (2017-18), Maria Gandin and Elena Marcucci (2018), Enrico Alfieri (2019), Anna Roncagalli (2020-21), Elia Manolino (2020-21), Gabriele De Francesco (2021-22), and Siria Carta (2021-22). Special thanks to the over 10,000 women and men of all eye colors who were curious and kind enough to take part in these studies.

## Notes

https://bit.ly/3Sq3Kf9

### Competing Interest Statement

The authors have declared no competing interest.

